# Atomic force microscopy of phase separation on ruptured, giant unilamellar vesicles

**DOI:** 10.1101/250944

**Authors:** Yanfei Jiang, Guy M. Genin, Kenneth M. Pryse, Elliot L. Elson

## Abstract

Giant unilamellar vesicles (GUVs) and supported lipid bilayers (SLBs) are synthetic model systems widely used in biophysical studies of lipid membranes. Phase separation behaviors of lipid species in these two model systems differ due to the lipid-substrate interactions that are present only for SLBs. Therefore, GUVs are believed to resemble natural cell membranes more closely, and a very large body of literature focuses on applying nano-characterization techniques to quantify phase separation on GUVs. However, one important technique, atomic force microscopy (AFM), has not yet been used successfully to study phase separation on GUVs. In the present study, we report that in binary systems, certain phase domains on GUVs retain their original shapes and patterns after the GUVs rupture on glass surfaces. This enabled AFM experiments on phase domains from binary GUVs containing 1,2-dilauroyl-sn-glycero-3-phosphocholine (DLPC) and either 1,2-dipalmitoyl-sn-glycero-3-phosphocholine (DPPC) or 1,2-distearoyl-sn-glycero-3-phosphocholine (DSPC). These DLPC/DSPC and DLPC/DPPC GUVs both presented two different gel phases, one of which (bright phase) included a relatively high concentration of DiI-C_20_ but excluded Bodipy-HPC, and the other of which (dark phase) excluded both probes. The bright phases are of interest because they seem to stabilize dark phases against coalescence. Results suggested that the gel phases labeled by DiI-C_20_ in the DLPC/DSPC membrane, which surround the dark gel phase, is an extra layer of membrane, indicating a highly curved structure that might stabilize the interior dark domains. This phenomenon was not found in the DLPC/DPPC membrane. These results show the utility of AFM on collapsed GUVs, and suggest a possible mechanism for stabilization of lipid domains.

## INTRODUCTION

The heterogeneity of cell membranes is generally believed to play important roles in a variety of cellular processes, although very little is known regarding the underlying physical basis by which the membranes accomplish these functions^1^.A current and popular hypothesis is that cell membranes contain dynamic sub-micrometer “lipid raft” domains that are enriched in certain kinds of lipids and protein species and phase separated from the rest of the membrane^2^. Since the earliest studies of phase separation on membranes, binary mixtures of lipids having long saturated acyl chains together with lipids having short or unsaturated chains have proven to be important model systems^3,4^. Upon cooling these binary systems from a state in which the two lipid components are uniformly mixed in a fluid phase, two phases emerge. The lipid with the higher transition temperature appears as an ordered solid (gel) while the lipid with the lower transition temperature remains in a disordered liquid state. Ternary mixtures containing the above two lipid species and cholesterol or other sterols can allow the emergence of a new phase: a liquid ordered phase. The liquid-ordered phase is believed to have lateral order similar to that in a liquid disordered phase, but configurational order of the hydrocarbon chains more similar to that in a gel phase due to the lipid-cholesterol interaction^5^. Lipid rafts are generally believed to be in the liquid-ordered state ^2,6,7^.

Motivated in part by the supposition that micron-sized domains in binary and ternary lipid systems serve as models for these nanoscopic lipid rafts, micron-sized lipid domains have long been studied in model membranes^6,8–11^. These can persist in a stable or metastable state over times sufficiently long to be relevant to physiologic processes. However, lipid rafts, if they exist, are nanoscopic. It is not clear whether simple lipid phase domains can persist for times long enough to be relevant physiologically and if so, what forces stabilize the domains at the nanoscopic range. This motivates a broad range of experimental and theoretical studies of model lipid membrane systems ^12–15^. Several potential factors have been proposed, including entropic force and mechanical repulsion caused by the difference of the intrinsic curvature of different phases^1,16,17^.

Several model systems are in widespread use to study membrane heterogeneity and phase separation. These include supported monolayers or bilayers, and synthetic vesicles, all of which can be synthesized with controlled compositions ^6,18^. Among these, giant unilamellar vesicles (GUVs) are particularly promising. First, compared to other vesicle models, their size range falls into that of mammalian cells (tens of micrometers). This is important because one driving force of phase separation relates to differences between the curvature of a membrane and the curvature that a lipid species would adopt in the absence of mechanical constraint ^12,13,17^. Second, compared to supported lipid monolayers and bilayers, lipid molecules diffuse more freely on vesicles due to the absence of lipid-substrate interactions. Indeed, phase separation behavior observed on supported bilayers differs from that observed on GUVs ^19^.

As reviewed elsewhere ^1^a broad range of characterization technologies have been applied to image micro- and nanodomains on model membranes. Phase separation can be visualized directly with either atomic force microscopy (AFM)^20,21^ or fluorescence microscopy ^8,9^. AFM has the advantages of nanoscale resolution and the detection of thickness differences amongst membrane phases. Fluorescence microscopy, which involves lipid probes with different affinities to the different phases, allows for color rendering of phase behavior over large regions and, in principle, for application of statistical fluorescence fluctuation technologies to characterize nanodomain dynamics ^6,8,9,22,23^.

However, each of these methods and models has limitations. Phase separation in GUVs has not been studied using AFM due not only to routine challenges associated with probing compliant structures ^24^ but also to additional challenges associated with thermal fluctuations^1^. Fluorescence fluctuation methods can reveal much about phase domain dynamics, but do not have the combination of spatial and temporal resolution needed to provide detail sufficient to image mechanisms that might contribute to stability of nanodomains^1^.

In the present study, we found that the patterns and shapes of gel domains in phase separated GUVs, as observed using fluorescence microscopy, persisted in the membrane adherent to a glass surface after the GUVs were ruptured. This enabled further AFM measurements on these domains. We applied this method to two binary systems: GUVs consisting of a mixture of 1,2-dilauroyl-sn-glycero-3-phosphocholine (DLPC) and either 1,2-dipalmitoyl-sn-glycero-3-phosphocholine (DPPC) or 1,2-distearoyl-sn-glycero-3-phosphocholine (DSPC). These two systems are unusual in that they present two different gel phases that appear to exist in equilibrium with one another. One of these gel phases recruits a high concentration of the fluorescent probe DiI-C_20_ but excludes the fluorescent probe Bodipy-HPC, while the other gel phase excludes both of these probes. The DiI labeled phase appears as an interphase enclosing the second gel phase and separating domains of the second gel phase both from each other and from the outside liquid disordered phase.

In the following, we present details of the technique for preserving phase behavior while collapsing GUVs onto glass slides, and apply this technique to characterize the ways that a DiI-labeled interphase might contribute to stabilizing the second gel phase. Results suggest that the DiI-labeled interphase in the DLPC/DSPC membrane exists as an extra layer of membrane in the ruptured GUVs. We propose that this is caused by mechanical effects associated with a highly curved structure of the DiI-labeled interphase. The extra layer of membrane was not found on the collapsed DLPC/DPPC membrane, suggesting that mechanical effects sufficient to block coalescence of microdomains do not require an out-of-plane membrane domain. We discuss these observations in the context of other models of curvature-driven membrane domain stabilization.

## MATERIAL AND METHODS

### Lipids and GUV preparation

GUVs were synthesized from lipids (Avanti Polar Lipids, Alabaster, AL), Bodipy-HPC fluorescent dye (Molecular Probes, Eugene, OR, number D3803), and DiI-C_20_ (Molecular Targeting Technologies, West Chester, PA). Lipids were stocked at 10 mM in chloroform, DiI-C_20_ in ethanol at 0.1 mM, and Bodipy-HPC in ethanol at 0.5 mM. GUVs were electroformed^10^. Briefly, 3 μl DSPC (or DPPC), 7 μl DLPC, 1 μl Bodipy-HPC and 1 μl DiI-C_20_ stock solutions were mixed and deposited onto two platinum wires by dragging the lipid solution along the wires back and forth until the chloroform and ethanol dried. The two wires were then placed into a homemade Teflon cylinder chamber filled with 300 mM sucrose solution. The chamber cap had two holes to hold the wires and separate them at a distance of 2 mm. The chamber was next placed upon a heating stage which maintained a temperature of 70°C. The wires were then connected to a function generator providing a 10 Hz/1.5 V square waveform for 1-2 hours.

### Fluorescence confocal microscopy

After the GUV solution cooled to room temperature, it was transferred from the Teflon chamber to an 8-well chamber slide (Lab-Tek) or glass cover slip (Fisher No. 0) with 300 μl distilled water or 300 mM glucose. Fluorescence confocal microscopy was performed on a LSM 510 ConfoCor 2 Combination System (Carl Zeiss, Germany) based on a Zeiss Axiovert 200M inverted microscope. A C-apochromatic 40X water immersion objective (numerical aperture 1.2) was used.

### AFM

Scanning was performed using an AFM (Asylum Research MFP-3D-BIO, Goleta, CA) in tapping mode using a cantilever with a resonant frequency of 10 kHz, a spring constant of 0.02 N/m, and a 4-sided pyramid tip with a radius of 40 nm (iDrive Magnetic Actuated Cantilever, Asylum Research, Goleta, CA). The AFM head was mounted on an Olympus X711 microscope. Fluorescence images were taken using a 60X Olympus oil objective. To image the lipid layers, a cover glass (Fisher No. 0) was first cleaned using Fisher lens cleaning paper and then placed onto the microscope. Next, 200 μl 300 mM glucose solution was pipetted onto the cover glass. 50 μl GUV solution was then transferred from the Teflon chamber to the glucose solution on the cover glass. After 15-30 minutes, the cover glass was gently rinsed using water. About 200-400 μl of water were left after the rinsing step. Finally, the AFM was mounted on the microscope stage for AFM measurements.

## RESULTS AND DISCUSSION

Upon cooling the GUVs, DSPC separated from DLPC to form two distinct gel phases (Figure 1). These two gel domains shared many similarities with patch and stripe domains reported previously and which we also observed (Figure 4) in DLPC/DPPC membrane systems ^11,21^. As reported by Li, et al^11^., the patch domains excluded both Bodipy-HPC (green) and Rh-DOPE/DiI-C_18_ (red) while the stripe domains attracted high concentrations of Rh-DOPE or DiI-C_18_ and low concentrations of Bodipy-HPC. The DLPC/DPPC patch and stripe domains differed from those observed on DLPC/DSPC GUVs: the red (DiI or Rh-DOPE) domains in the DLPC/DSPC GUVs are stripe-like and have smooth edges,^11^ while the red domains in the DLPC/DSPC GUVs were irregular with rough edges (Figure 1A). Therefore we term the dye-rich gel domains “bright domains” in the following, and distinguish them from the patch-like “dark domains.” Bright domains tended to form on the edges of dark domains in DLPC/DSPC GUVs, potentially affecting the latter both thermodynamically and kinetically.

**Figure 1.**
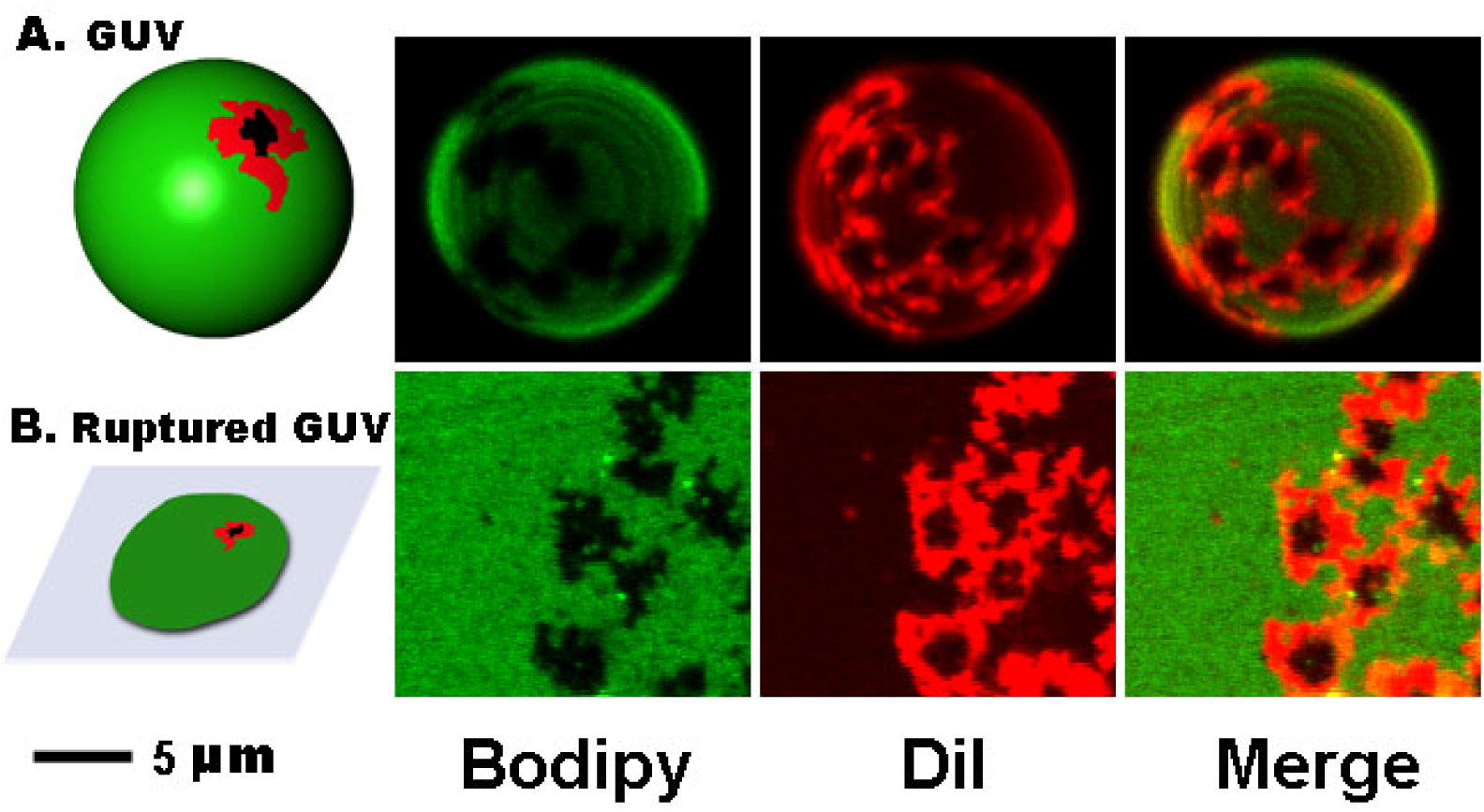
Confocal images of a DLPC/DSPC GUV and ruptured GUVs on a cover slip. GUVs are labeled with Bodipy-HPC (green) and DiI-C_20_ (red). GUVs are made of 30% DSPC and 70% DLPC.

The GUVs studied contained 300 mM sucrose solution and therefore sedimented to the glass slide when transferred to a lower density solution (300 mM glucose or water). There, they ruptured and spread over the glass surface. We found that the patterns and shapes of the gel domains in DLPC/DSPC GUVs were preserved after the GUV ruptured (Figure 1B). The bright domains remained around the edges of the dark domains and retained their irregular shapes.

We conducted AFM experiments and compared tapping mode scans with fluorescence images to study structural differences between these two different gel domains (Figure 2). Interpretation of the topologies obtained required first identifying the source of what appeared to be “nanodomains” in the liquid phase area in the AFM scans but that did not appear in the fluorescence images (Figure 2, B and C, Line 1). These “nanodomains” were several hundred nanometers in diameter and about 2 nm thicker than the surrounding membrane (Figure 2C, Line 1). They were also observed when the DLPC:DSPC ratio was decreased to 5:95, a concentration at which the system is expected to be in the liquid phase at 22°C ^25^ (supporting material). Further, they did not melt or change shape when the temperature was increased to 35°C (Figure S1). These results suggest that these “domains” are not DSPC but rather DLPC bilayer “islands” embedded in a surrounding monolayer membrane. Although one might expect that such bilayers embedded within monolayers might appear as nonuniformities in the fluorescence intensity maps, we note that these bilayer “islands” have a size that is near the optical resolution limit and are thus hard to detect using the regular fluorescence microscope installed on our AFM system. Indeed, confocal images of the liquid phase area with better resolution show that the fluorescence in the liquid phase region of collapsed GUVs is nonuniform (supporting material, Figure S2).

**Figure 2.**
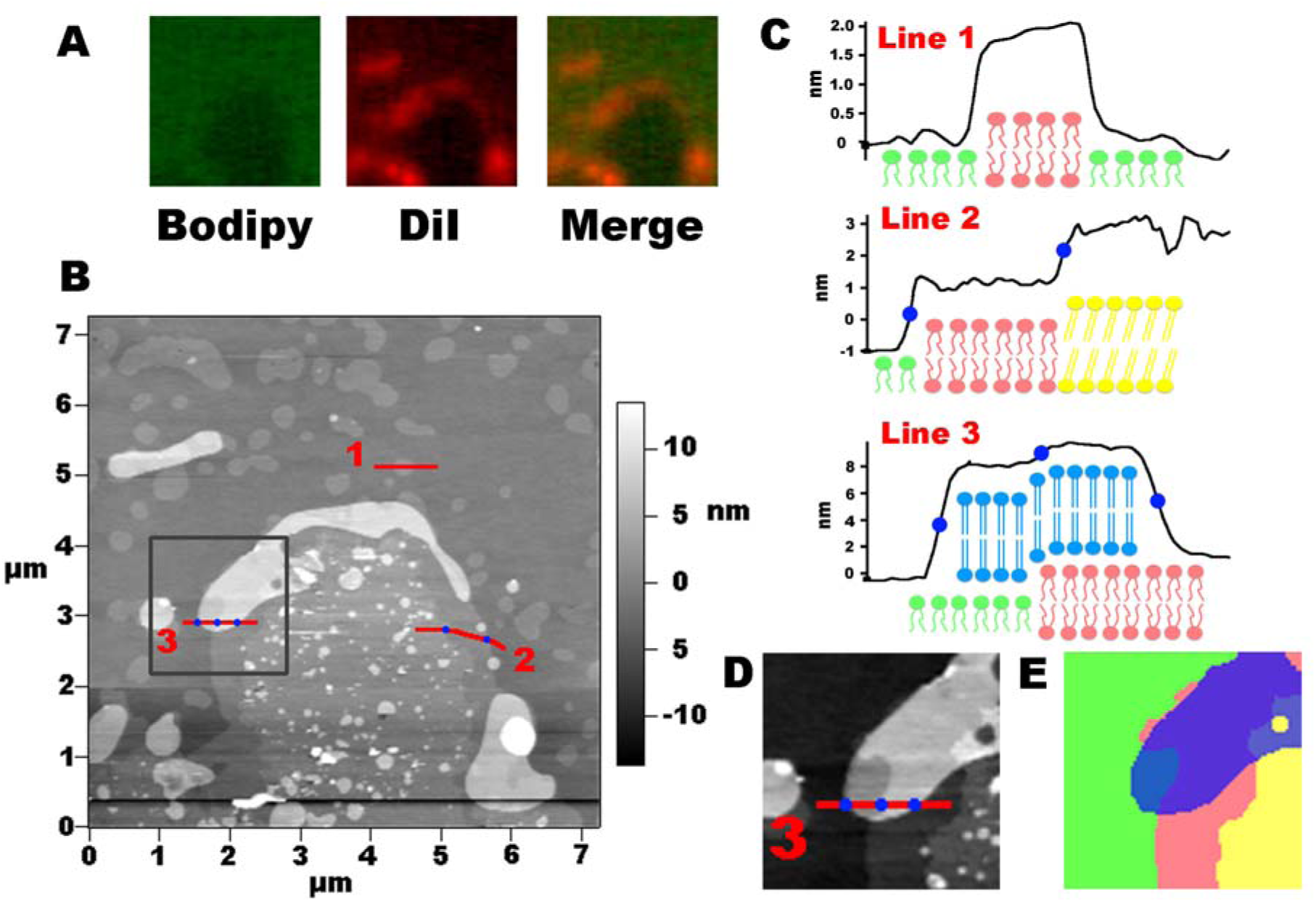
AFM measurements on ruptured GUVs. (A) Fluorescence image of domains on which the AFM scanning was performed. Images were taken using a standard fluorescence microscope instead of the confocal microscope used for **Figure** 1, resulting in a lower resolution. (B) Height profiles from the domains shown in panel A. (C) Height profiles along the three lines shown in panel B. The origins of the curves (left) correspond to the numbered ends of lines shown in panel B. Cartoons of colored lipid molecules correspond to the hypothesized stacking of lipid layers (green: DLPC monolayer; pink: DLPC bilayer; yellow: DSPC bilayer, dark domain; blue: DSPC bilayer, bright domain). (D) Magnified image from panel B. The contrast has been changed for better illustration of the topology. (E) Color rendering of the topology in panel D. Colors match those of panel C.

With this information in hand, we proceeded to characterize the unusual topologies of the two gel phases. The height profile of a line drawn from the monolayer liquid phase to the inside a dark domain revealed a stair topology of two steps (Line 2 in Figure 2 B and C). The first step was about 2 nm high and the second step was about 1.8 nm high. As discussed above, the first step corresponded to a progression from the DLPC monolayer to a bilayer. The second step therefore was interpreted as a progression from a DLPC bilayer to a DSPC bilayer. The height difference of 1.8 nm is consistent with published thickness differences between DLPC and DSPC bilayers studies via AFM ^20^.

In contrast to the dark domains that were connected directly to the DLPC liquid phase, the bright domains were found to consist of an extra layer of membrane residing atop the remainder of the membrane. This surprising conclusion was reached based upon three observations. First, the height difference between the DLPC monolayer and the bright domains was around 8 nm, which was too thick to be explained by a new phase of a bilayer (Figure 2, B and C, Line 3). Second, the bright domains could be knocked off of the dark domain progressively using and a scanning AFM tip in tapping mode (Figure 3A). The loss of the bright domains was much faster with contact mode scanning (data not shown). This was confirmed by fluorescence microscopy before and after experiments designed to knock off the bright domains (Figure 3B). Third, the height measured via AFM changed with that of the lipids supporting the bright domains. For example, as a bright domain extended from a DLPC monolayer to a DLPC bilayer (second blue point in Figure 2, B and C), its thickness remained at 8 nm (third blue point) while the overall height rose 2 nm. This further suggests that the bright domains could retain their cohesion over height steps even greater than those found at the interface between DSPC and DLPC bilayers, and that, although they existed in the gel phase, they were sufficiently compliant in shear to follow the contour of the lipids beneath them.

**Figure 3.**
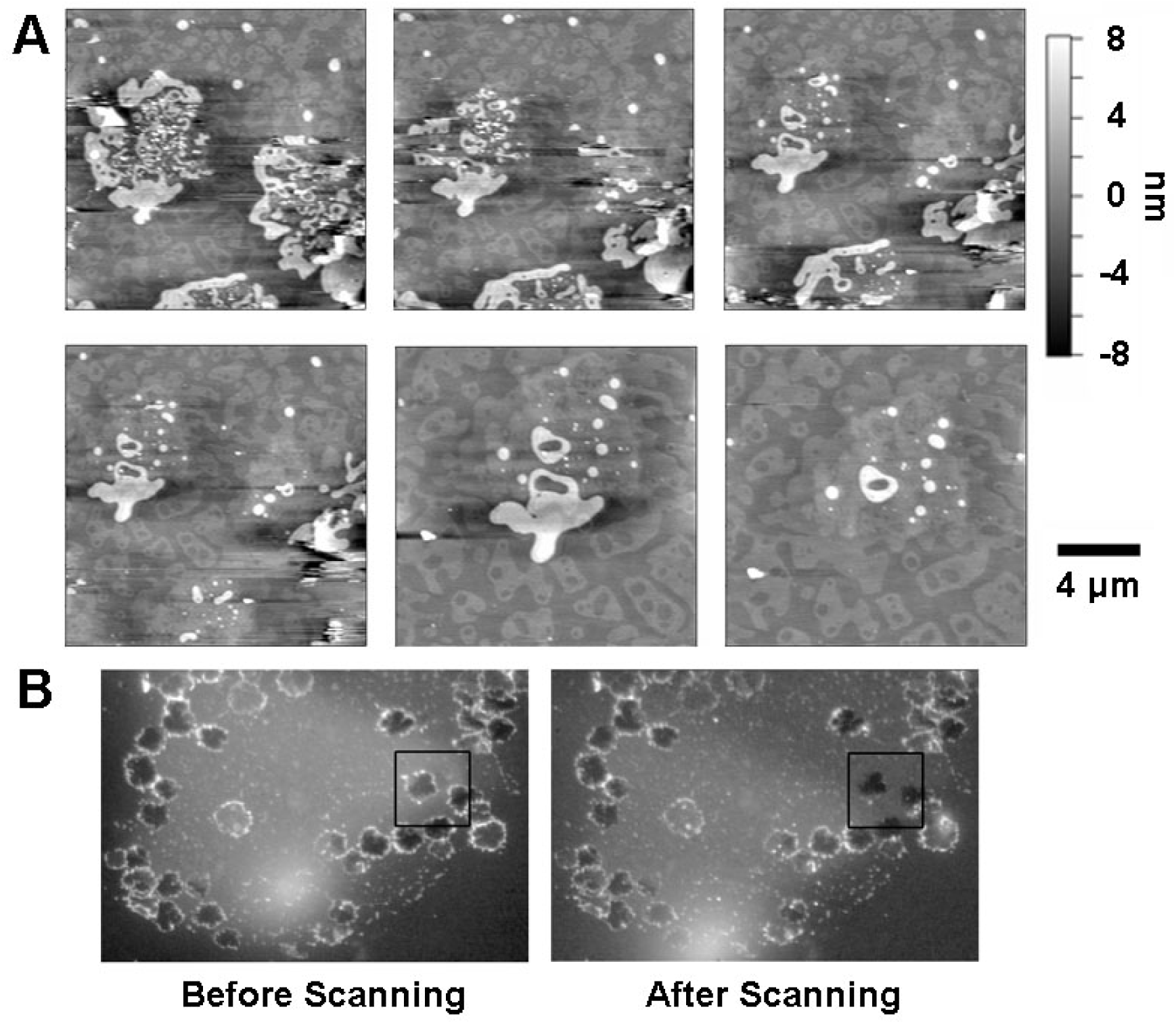
Bright domains on DLPC/DSPC collapsed GUVs could be knocked off by an AFMtip. (A) Height images from a continuous scanning. The scale bar is for the first four images. For the last two images, showing the domain that appears on the left side in the first four images, the scale bar indicates 6 micrometers. (B) Fluorescence images (DiI-C_20_) of the domains before and after the AFM scanning. The scanning area is indicated by the black square.

The thickness of the bright domains (8 nm) in our study was large but consistent with thickness measurements reported in the literature for bilayers supported atop other bilayers. We note a broad range of membrane thicknesses reported from AFM experiments in the literature, and propose that they can be reconciled as follows. AFM measurements are reported for both membrane layers supported directly by mica substrate and for additional membrane layers atop such membrane layers. We found that the thicknesses reported for second layers of membrane are between 6-9 nm and much thicker than those reported for a first layer, which are between 3-5 nm ^20,26,27^. Although the reasons for this are not clear, we suspect different water layer thicknesses between membrane/membrane and membrane/substrate interfaces, and note that our measurements are consistent with the former.

As discussed above, the bright domains have a high concentration of DiI, posing a question of whether the edge domains are simply aggregated DiI around the dark domains surrounding DSPC gel. To answer this question, GUVs were made without DiI and only labeled with Bodipy-HPC. AFM experiments performed on these ruptured GUVs showed that the 8 nm edge features were still present around the dark domains (Figure 4). This suggests that the edge domains are neither aggregations of DiI nor artifactual phases induced by DiI. Instead, the two-gel-phase separation appears to be intrinsic to the DLPC/DSPC membrane.

**Figure 4.**
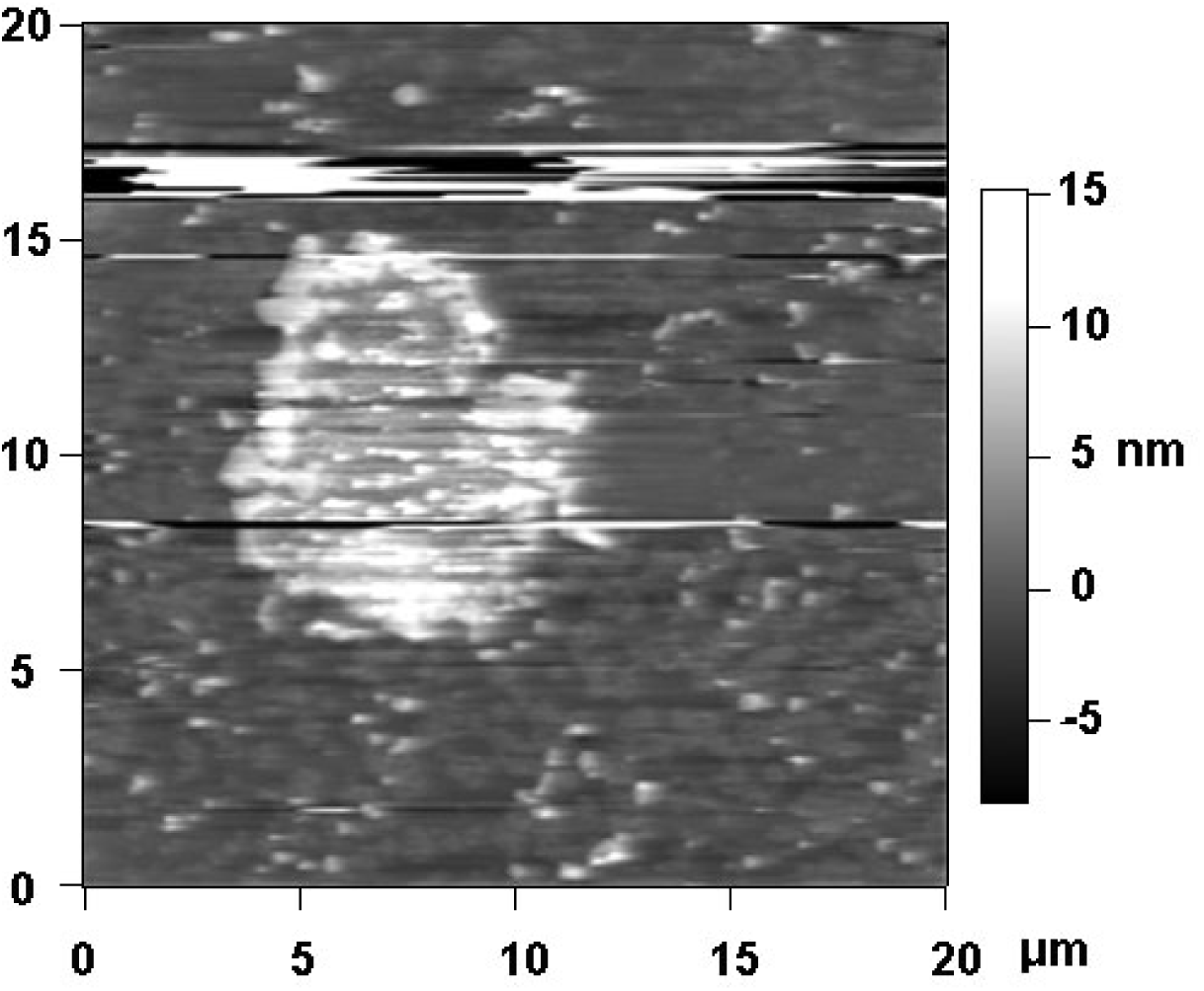
Height image of a dark domain on a ruptured DLPC/DSPC GUV membrane. TheGUV was labeled only with Bodipy-HPC. The image shows that the second gel phase was not dependent upon DiI-C_20_.

This conclusion extends that of Loura, et al., ^22^ who performed fluorescence lifetime and Förster resonance energy transfer measurements on DiI in a DLPC/DSPC membrane. Their results indicate the segregation of DiI into the gel/fluid interface, but they were not able to test this hypothesis on the small vesicles they employed. By showing interfacial domains with a high concentration of DiI, our confocal images of GUVs provide direct support of their conjecture that DiI might segregate into the gel/fluid interface. However, the dimension of interfacial domains we have observed is wider than a single molecular interface layer between the phases.

The application of AFM to collapsed GUVs enabled this to be explored more deeply and showed that instead of being just an single boundary layer of DiI, there is in fact a distinctgel phase that is rich in DiI.

Although this is the first report of two gel phases coexisting in the DLPC/DSPC system, a similar phenomenon has been reported in a DLPC/DPPC system in both GUVs ^11^ and supported lipid bilayers^21^. However, several differences are evident between these two membrane systems. As discussed above, the bright domains in DLPC/DPPC GUVs are stripe-like and have smooth edges, with the width of the bright domains homogeneous along the stripes. Bright domains are readily observed for DLPC/DPPC GUVs with a branching/turning pattern having a characteristic turning angle of 60°.^21^ In contrast to this highly ordered pattern, bright domains in DLPC/DSPC GUVs were more irregular and their edges were rougher. Combined AFM and fluorescence microscopy experiments showed that ruptured DLPC/DPPC GUVs also preserved their domain patterns following rupture and flattening, with smooth edges of constant width evident both before and after rupturing of GUVs (Figure 5). Scans analogous to those performed above showed that, in contrast to the DLPC/DSPC system, the dark and bright domains in a DLPC/DPPC membrane show no height difference (Figure 5B), and no evidence of the bright layer residing atop other membrane components.

**Figure 5.**
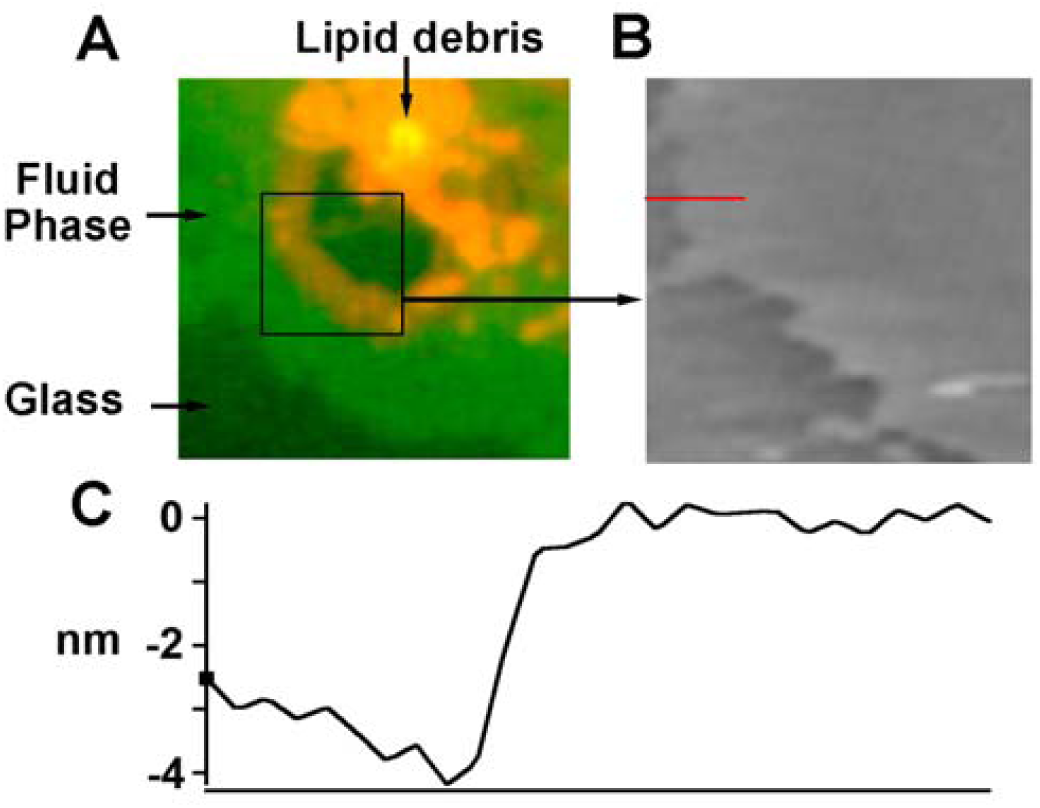
AFM of ruptured DLPC/DPPC GUVs. (A) Fluorescence image; (B) AFM height image; (C) Height profile along the red line in Figure B.

The fact that the DSPC and DPPC GUVs have different morphologies is not surprising given that their spontaneous curvatures have been estimated to be different by approximately a factor of two^28^. However, a question that we could not answer is whether the bright domain exists as an external layer on DLPC/DSPC GUVs or whether this can occur only on collapsed GUVs. In other words, were the DSPC lipid molecules pushed out of the GUVs during phase separation, or did this occur during rupture? Differences in spontaneous curvature and area changes associated with phase demixing are potential sources of strain energy that could drive either buckling or ejection of a gel phase. Alternatively, a structural instability in a bright phase might stay connected to the mother membrane before the GUV ruptures, but become disconnected and form a second layer of membrane as the GUV ruptures and spreads over the substrate. Our AFM results show that the width of the bright domain is usually below 1 micrometer on the ruptured membrane, so the radius of budding in GUVs would be close to or below the optical resolution limit (around 250 nm), making such bulges difficult to observe optically. Therefore, it is difficult to delineate the topology in the unruptured GUVs using current optical techniuqes.

Bulges, buckles, or ejected lipid could all lead to high local curvature capable of inducing a repulsive interaction between domains.^17^ This would have consequences important to the study of the membrane domain stability because a repulsive interaction of this character could increase the energetic barrier to domain coalescence and thereby serve to slow coarsening of domains.

This provides motivation for extrapolating our findings from the microscopic level to the nanoscopic level and for investigating how the bright interphase impacts the stability of nanodomains. Further studies of ruptured GUVs using techniques such as high resolution secondary ion mass spectroscopy^29^ and coherent anti-strokes Raman spectroscopy^11^ might shed additional light on these questions.

## CONCLUSION

Collapsing both DLPC/DPPC and DLPC/DSPC GUVs onto glass slides enables the application of AFM to the study of phase behavior in a way that can be related to behavior of GUVs. Although questions arise about the disconnection of bright domains from the lipid bilayer, the overall shape and patterns of domains are preserved after GUV rupture. Our findings provides a basis for further AFM investigations of the nanoscopic phase separation in GUVs, which has proven difficult to study due to the lack of powerful techniques that can break though the optical limit in a highly diffusive environment. Our results show that two distinct gel phases can exist simultaneously and interact on GUVs. The so-called bright domains can exist as an extra layer of membrane on ruptured GUVs possibly indicating a highly curved structure on unruptured GUVs, which might have important implications for nanodomain stability.

## ASSOCIATED CONTENT

### Funding Sources

This work was funded in part by the National Institutes of Health through grant R01GM084200, and through the National Science Foundation through grant CMMI 0826518.

### Supporting Information

Supporting Information Available: Figure S1 and S2. These materials are available free of charge via internet at http://pubs.acs.org

## ABREVIATIONS

DLPC: 1,2-dilauroyl-sn-glycero-3-phosphocholine
DPPC: 1,2-dipalmitoyl-sn-glycero-3-phosphocholine
DSPC: 1,2-distearoyl-sn-glycero-3-phosphocholine
GUV: giant unilamellar vesicle
SLB: supported lipid bilayer
AFM: atomic force microscopy

## REFERENCES

Elson E. L., Fried E., Dolbow J. E. & Genin G. M. Phase separation in biological membranes: integration of theory and experiment. Annu. Rev. Biophys. 39, 207–226 (2010).

Pike L. J. Rafts defined: a report on the Keystone Symposium on Lipid Rafts and Cell Function. J. Lipid Res. 47, 1597–1598 (2006).

Hinz H. J. & Sturtevant J. M. Calorimetric studies of dilute aqueous suspensions of bilayers formed from synthetic L- -lecithins. J. Biol. Chem. 247, 6071–6075 (1972).

Melchior, D. L., Morowitz, H. J., Sturtevant, J. M. & Tsong, T. Y. Characterization of the plasma membrane of Mycoplasma laidlawii. VII. Phase transitions of membrane lipids. Biochim. Biophys. Acta 219, 114–122 (1970).

Harder, T. & Sangani, D. Plasma membrane rafts engaged in T cell signalling: new developments in an old concept. Cell Commun. Signal. Ccs 7, 21 (2009).

Simons, K. & Vaz, W. L. C. Model systems, lipid rafts, and cell membranes. Annu. Rev. Biophys. Biomol. Struct. 33, 269–295 (2004).

Mouritsen, O. G. Theoretical models of phospholipid phase transitions. Chem. Phys. Lipids 57, 179–194 (1991).

Korlach, J., Schwille, P., Webb, W. W. & Feigenson, G. W. Characterization of lipid bilayer phases by confocal microscopy and fluorescence correlation spectroscopy. Proc. Natl. Acad. Sci. 96, 8461–8466 (1999).

Feigenson G. W. & Buboltz J. T. Ternary Phase Diagram of Dipalmitoyl-PC/Dilauroyl-PC/Cholesterol: Nanoscopic Domain Formation Driven by Cholesterol. Biophys. J. 80, 2775–2788 (2001).

Morales-Penningston N. F. et al. GUV preparation and imaging: minimizing artifacts. Biochim. Biophys. Acta 1798, 1324–1332 (2010).

Li L. & Cheng J.-X. Coexisting stripe- and patch-shaped domains in giant unilamellar vesicles. Biochemistry (Mosc.) 45, 11819–11826 (2006).

Embar A., Dolbow J. & Fried E. Microdomain evolution on giant unilamellar vesicles. Biomech. Model. Mechanobiol. 12, 597–615 (2013).

Maleki M. & Fried E. Multidomain and ground state configurations of two-phase vesicles. J. R. Soc. Interface R. Soc. 10, 20130112 (2013).

Brewster R. & Safran S. A. Line active hybrid lipids determine domain size in phase separation of saturated and unsaturated lipids. Biophys. J. 98, L21–23 (2010).

Palmieri B. & Safran S. A. Hybrid lipids increase the probability of fluctuating nanodomains in mixed membranes. Langmuir Acs J. Surfaces Colloids 29, 5246–5261 (2013).

Frolov V. A. J., Chizmadzhev Y. A., Cohen F. S. & Zimmerberg J. ‘Entropic traps’ in the kinetics of phase separation in multicomponent membranes stabilize nanodomains. Biophys. J. 91, 189–205 (2006).

Ursell T. S., Klug W. S. & Phillips R. Morphology and interaction between lipid domains. Proc. Natl. Acad. Sci. 106, 13301–13306 (2009).

Ulrich, A. S. Biophysical aspects of using liposomes as delivery vehicles. Biosci. Rep. 22, 129–150 (2002).

Tokumasu, F., Jin, A. J., Feigenson, G. W. & Dvorak, J. A. Nanoscopic lipid domain dynamics revealed by atomic force microscopy. Biophys. J. 84, 2609–2618 (2003).

Lin, W.-C., Blanchette, C. D., Ratto, T. V. & Longo, M. L. Lipid asymmetry in DLPC/DSPC-supported lipid bilayers: a combined AFM and fluorescence microscopy study. Biophys. J. 90, 228–237 (2006).

Bernchou, U., Midtiby, H., Ipsen, J. H. & Simonsen, A. C. Correlation between the ripple phase and stripe domains in membranes. Biochim. Biophys. Acta 1808, 2849–2858 (2011).

Loura, L. M., Fedorov, A. & Prieto, M. Partition of membrane probes in a gel/fluid two-component lipid system: a fluorescence resonance energy transfer study. Biochim. Biophys. Acta 1467, 101–112 (2000).

Pryse, K. M. et al.Confidence intervals for concentration and brightness from fluorescence fluctuation measurements. Biophys. J. 103, 898–906 (2012).

Tsukruk, V. V. & Singamaneni, S. Scanning Probe Microscopy of Soft Matter. (2011).

Mabrey, S. & Sturtevant, J. M. Investigation of phase transitions of lipids and lipid mixtures by sensitivity differential scanning calorimetry. Proc. Natl. Acad. Sci. 73, 3862–3866 (1976).

Leidy, C., Kaasgaard, T., Crowe, J. H., Mouritsen, O. G. & Jørgensen, K. Ripples and the formation of anisotropic lipid domains: imaging two-component supported double bilayers by atomic force microscopy. Biophys. J. 83, 2625–2633 (2002).

Simonsen, A. C. & Bagatolli, L. A. Structure of spin-coated lipid films and domain formation in supported membranes formed by hydration. Langmuir Acs J. Surfaces Colloids 20, 9720–9728 (2004).

Kollmitzer, B., Heftberger, P., Rappolt, M. & Pabst, G. Monolayer Spontaneous Curvature of Raft-Forming Membrane Lipids. (2013). at <http://arxiv.org/abs/1307.0367>

Kraft, M. L., Weber, P. K., Longo, M. L., Hutcheon, I. D. & Boxer, S. G. Phase Separation of Lipid Membranes Analyzed with High-Resolution Secondary Ion Mass Spectrometry. Science 313, 1948–1951 (2006).

